# Versatile and Robust method for Antibody Conjugation to Nanoparticles with High Targeting Efficiency

**DOI:** 10.1101/2021.09.29.462399

**Authors:** Indra Van Zundert, Maria Bravo, Olivier Deschaume, Pierre Cybulski, Carmen Bartic, Johan Hofkens, Hiroshi Uji-i, Beatrice Fortuni, Susana Rocha

## Abstract

The application of antibodies in nanomedicine is now standard practice in research since it represents an innovative approach to deliver chemotherapy agents selectively to tumours. The variety of targets or markers that are overexpressed in different types of cancers results in a high demand for antibody conjugated nanoparticles which are versatile and easily customizable. Considering upscaling, the synthesis of antibody conjugated nanoparticles should be simple and highly reproducible. Here, we developed a facile coating strategy to produce antibody conjugated nanoparticles using ‘click chemistry’ and further evaluated their selectivity towards cancer cells expressing different markers. Our approach was consistently repeated for the conjugation of antibodies against CD44 and EGFR, which are prominent cancer cell markers. The functionalized particles presented excellent cell specificity towards CD44 and EGFR overexpressing cells, respectively. Our results indicated that the developed coating method is reproducible, versatile, non-toxic, and can be used for particle functionalization with different antibodies. This grafting strategy can be applied to a wide range of nanoparticles and will contribute to the development of future targeted drug delivery systems.

## 1. Introduction

Despite the numerous advances in treatment options, cancer remains a leading cause of mortality worldwide. Existing methods bear limitations and complications, such as incomplete removal of the tumour and severe side-effects. Therefore a combination of treatments is often required to reach the desired results [1–3]. The urge to develop more effective therapies gave rise to intensive research in delivery of chemotherapeutics using nanoparticles. Engineered nanoparticles have shown to serve as excellent drug nano-carriers. Among the advantages of using nanoparticles are their higher drug loading capacity, the protection of the drugs against degradation during blood circulation and the possibility to easily add other functionalities. As the size of particles can be tailored, nanoparticles between 20 and 200 nm can take advantage from the enhanced permeability and retention (EPR) effect, to passively accumulate near the tumour because of abnormal blood vessel architecture [4,5]. However, over the last years, increasing debate on the EPR effect has emerged, raising doubts about its reliability and applicability [6–8]. To this end, active targeting of nanoparticles to cancer cells is preferred.

Over the years, a wide range of nanoparticles have been engineered and several strategies have been developed to promote nanoparticle internalization into specific cells. Often, nanoparticles are functionalized with ligands that recognize overexpressed receptors or markers present on the cancer cell membrane [5,9,10]. In doing so, they facilitate specific accumulation of the drug in cancer cells [11]. Folic acid or transferrin conjugated nanoparticles are popular examples of such drug delivery systems (DDSs), as they bind to folate and transferrin receptors, respectively, overexpressed in certain cancers [10,12,13]. Typically, one nanoparticle is designed against a particular receptor or marker, hence targeting a specific cancer. However, patients with the same type of cancer can overexpress different markers. For instance, overexpression of the estrogen receptor (ER) is linked to a hormone sensitive form of breast cancer (ER+), while HER2 is overexpressed in an aggressive and fast-growing type of breast cancer (HER2+)[14–16]. Given the variety in potential targets, there is a continuous search for simple methods to customize nanoparticles, turning them into versatile nano-carriers. To this end, antibodies have proven to be a promising strategy as they can be developed against most of the existing markers. The success of antibodies in targeting was already proved with the arrival of antibody-drug conjugates, that have emerged as powerful therapeutic agents in cancer therapies and are extensively used in the clinic today [17–19]. After the advances in antibody-drug conjugates, conjugation of antibodies to nanoparticles yields great therapeutic potential [20,21].

Conjugation of antibodies to nanoparticles can be achieved via different strategies i.e. ionic adsorption (non-covalent attachment) [22], covalent binding (including carbodiimide chemistry [23], maleimide chemistry [24] and click-chemistry [25]) or using adapter molecules such as biotin [26]. In ionic adsorption, antibodies are linked to the nanoparticles via electrostatic interactions [27], leading to poor reproducibility and low stability [28]. Alternatively, adapter molecules, such as the avidin-biotin couple, can be implemented, but this interaction is influenced by the pH, affecting nanoparticle stability in more acidic conditions as found in the tumour microenvironment [29].

Covalent attachment is achieved by functionalizing the nanoparticle surface with functional groups (e.g. amine, carboxylic, maleimide,…) which can react with the amino acid side chain of the antibody by standard conjugation methods. Covalent attachment of antibodies is generally preferred, provided that an appropriate approach is used. For instance, while EDC/NHS coupling is a common method for covalent attachment [30], it can result in oligomerization of antibody molecules [31]. Furthermore, some conjugation methods require the use of catalysts, often metals, that can lead to increased toxicity of the nanoparticles if not fully removed from the solution [32,33]. Therefore, when using covalent conjugation, a catalyst-free approach and vast optimization are important. Despite the progress in nanoparticle functionalization, there is still an urge for simple and reproducible strategies to conjugate antibodies to nanoparticles, enabling the development of versatile DDSs which can easily be customized for selectivity towards different cancer markers.

In this work, we propose a simple and reproducible coating strategy for antibody conjugation to nanoparticles. To avoid the use of catalysts, we developed an approach based on copper free click chemistry (Figure 1). Briefly, antibodies were labelled with a dibenzocyclooctyne (DBCO) moiety, while an azide (N_3_) group was attached to the nanoparticles. The independent activation of the antibody and nanoparticles reduces the possibility of oligomerization of the antibody or aggregation of the particles. We prove the versatile nature of our method by creating two different types of particles, either conjugated to an anti-Cluster of Differentiation 44 (anti-CD44) or anti-Epidermal Growth Factor Receptor (anti-EGFR) antibody, targeting CD44 or EGFR overexpressing cells, respectively. CD44 and EGFR are surface receptors that manifest themselves as important biomarkers in cancer [34,35]. In this report, mesoporous silica nanoparticles (MSNPs) were used as a model application for our coating strategy. In recent years, MSNPs have been pointed out as extremely promising tools in cancer research given their high biocompatibility, chemical stability, high drug loading and releasing capacity, straight-forward functionalization, and low-cost, scalable fabrication [36–38]. For these reasons, MSNPs were chosen as a study model of DDSs for further surface modifications. Nevertheless, we foresee that the coating strategy here presented can be easily applied to a wide range of nano-carriers besides MSNPs, as it only requires the presence of amine groups on the surface of the nanoparticle. This simple conjugation strategy will contribute to the up-scaling of antibody conjugated nanoparticles and to the future developments of targeted nanoparticles with multiple functionalizations.

**Figure 1:**
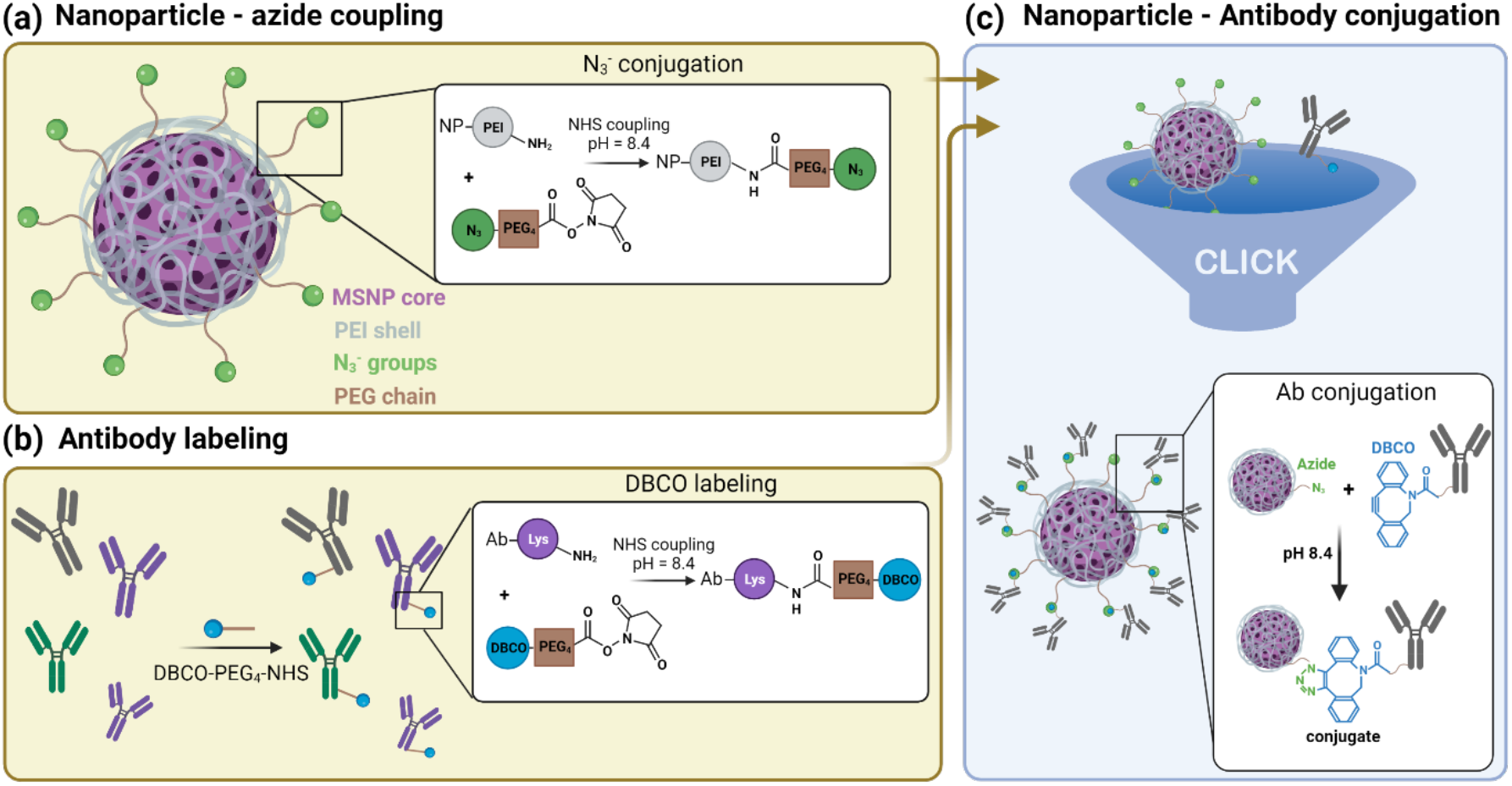
Schematic representation of the antibody-conjugated nanoparticle preparation: (a) Mesoporous silica nanoparticles (MSNPs) loaded with the drug (Doxorubicin) and coated with a polyethyleneimine (PEI) layer (gray) are grafted with an Azide moiety (N_3_, green) using NHS ester coupling. (b) The lysine residues (purple) of different antibodies were labelled with a DBCO moiety (blue) via NHS ester coupling. (c) Click chemistry reaction of the Azide functionalized nanoparticle with the DBCO labelled antibodies resulting in the final antibody conjugated nanoparticle.

## 2. Materials and Methods

### Materials

Anti-human EGFR (αEGFR, monoclonal, cat. BE0278) and anti-human CD44 (αCD44, monoclonal, hermes-1, cat. BE0039) were purchased from Bio X Cell (Lebanon). Tetraethyl orthosilicate (TEOS, 98%), cetyltrimethylammonium chloride solution (CTAC, 25% in H_2_O), triethanolamine (TEA, 99%), hydrochloric acid (HCl, 1 N), Rhodamine B basic violet 10 (RhoB, 93%), Fluorescein isothiocyanate (FITC), (3-Aminopropyl)triethoxysilane (APTES), polyethyleneimine solution (PEI, 50% w/v in H2O), Triton X-100 (0.1%), N-(3-dimethylaminopropyl)-N’-ethyl-carbodiimide (EDC, 97%), Dimethyl sulfoxide (DMSO, >99,5%) and 3D Petri Dish ® - Microtissues were purchased from Sigma Aldrich (Germany). Dulbecco’s modified eagle medium (DMEM), LysoTracker™ Deep Red, DiR lipophilic dye, Gentamicin, Dulbecco’s phosphate buffered saline (PBS, no calcium, no magnesium), Formaldehyde (4% in PBS), trypsin-EDTA (0.5%), Hank’s balanced salt solution (HBSS, no phenol red), Ethanol (absolute, 99.9%), donkey-anti-rat IgG-Alexa Fluor 488 (cat. A-21208), and Zeba™ Spin Desalting Columns (40K MWCO, 0.5mL) were purchased form ThermoFisher Scientific (United States). Atto488 and Atto565 NHS ester conjugate were purchased from Atto-TEC GmbH (Germany). DBCO-PEG4-NHS-ester and N_3_-PEG_4_-NHS-ester were purchased from Click Chemistry Tools (United States). All the chemicals were used without further purifications.

### CD44 and EGFR antibody labeling

The antibodies were functionalized with DBCO-PEG_4_-NHS ester for conjugation onto the nanoparticles via copper free click chemistry. In addition, a fluorescent label, Atto565-NHS ester or Atto488-NHS ester, was sometimes grafted onto the antibody for further investigations via fluorescence microscopy. The labelling of 500 µg antibody was carried out in 50 mM borate buffer pH 8.5 (antibody concentration 1 mg/mL). Five molar equivalents of DBCO-PEG_4_-NHS ester and two molar equivalents of atto570-NHS ester (when needed) were added to the reaction, according to Eggermont and Hammink *et al*. [39]. After 6 h under magnetic stirring at room temperature, the dual labelled antibodies were purified over a 0.5 mL 40K Zeba desalting column to remove residual free atto570-NHS ester and DBCO-PEG_4_-NHS ester molecules.

### Synthesis MSNPs

The MSNPs were synthetized by the biphase stratification method reported by Shen et al. [40]. In short, 0.18 g of TEA was mixed with a solution of 24 mL CTAC and 36 mL of Milli-Q water. This mixture was heated to 60°C under magnetic stirring for 1 h. Next, 20 mL of TEOS (20 v/v % in octadecene) was slowly added with a syringe and the reaction was kept proceeding overnight. When fluorescein (FITC) was linked to the MSNPs matrix (FITC encapsulation), 16.6 mg of FITC was dissolved in 10 mL 99,8% ethanol and 400 μL of (3-Aminopropyl)triethoxysilane (APTES). This mixture was stirred for 2 h under inert atmosphere to couple FITC to the aminosilane. After 2 h, the solution was added together with the TEOS. Next, the reaction was cooled down to room temperature and the nanoparticles were washed with a solution of HCl 1.1 M in water/ethanol (v/v = 1.25:10) with centrifugation-dispersion-sonication cycles to remove CTAC from the pores. Subsequently, the nanoparticles were washed two times with milli-Q water in order to neutralize the pH.

### MSNP dye/drug loading

The pores of the MSNPs were loaded with doxorubicin (Dox) or rhodamine B (RhoB) for cytotoxicity experiments and fluorescence imaging, respectively. Loading of RhoB was performed in milli-Q water under magnetic stirring for 3 h. For Dox loading, MSNPs were first dispersed in phosphate buffer (pH 9) to maximize the loading efficiency. To avoid Dox aggregation, the solution containing Dox and MSNPs was sonicated for 10 minutes. Next, the solution was stirred for 24 h at 400 rpm. After loading (of Dox or RhoB), the solution was centrifuged and the supernatant was replaced with Milli-Q water and the Dox- and RhoB-loaded nanoparticles were re-suspended (MSNPs_Dox and MSNPs_RhoB, respectively). The supernatants of all the centrifugation steps were collected and measured with a spectrometer in order to quantify the Dox loaded inside the MSNPs. We estimated a Dox concentration in the MSNPs of approximately 50 µM.

### MSNP functionalization

To coat the nanoparticles with a PEI layer, a 0.75% PEI solution (in Milli-Q water), adjusted to pH 7 (with 37% HCl) was added to the dye -or drug loaded MSNPs (1:1 ratio) in a plastic vial. This mixture was magnetically stirred for 3 h, yielding PEI-coated MSNPs. To facilitate the azide conjugation, PEI-coated particles were dispersed in borate buffer (pH 8.5). To conjugate the NHS ester-PEG_4_-N_3_ linker to the PEI amine groups, an NHS ester reaction was used. In detail, 1.5 mg NHS ester-PEG_4_-N_3_ (in DMSO) were added to 500 µL of PEI-MSNPs (10 mg/mL) in a dropwise manner and magnetically stirred for 4 h. After the reaction, the nanoparticles were centrifuged and re-dispersed in borate buffer (N_3_-PEI-MSNPs). To conjugate the CD44 antibody via a copper free click reaction, 100 µg labelled CD44 antibody (50 µL of 1 mg/mL labelled antibody solution in borate buffer) were added to 450 µL of N_3_-PEI-MSNPs (10 mg/mL). The reaction was stirred for 6 h at room temperature. After the reaction, nanoparticles (Ab-PEI-MSNPs) were centrifuged at low speed (700 RPM) and re-dispersed in Milli-Q water.

### Ab-PEI-MSNPs characterization

The synthesized nanoparticles were characterized by confocal fluorescence microscopy, scanning electron microscopy (SEM) and atomic force microscopy (AFM). For the fluorescence microscopy experiments, 60 µL Ab-PEI-MNSP solution was pipetted in a Coverwell™ perfusion chamber placed onto a #1 coverglass. After 30 min, when some nanoparticles have sedimented, the sample was imaged with a Leica TCS SP8 mini microscope. For SEM measurements, nanoparticles were drop casted on Indium-Tin Oxide coated glass and dried. Next, the glass was coated with Au/Pd for 20s. Nanoparticles were visualized using a FEI Quanta 250 FEG Scanning Electron Microscope. Zeta potential measurements were carried out on a Malvern Zetasizer system. For AFM characterization, an Agilent 5500 AFM with MAC III controller was used for morphological imaging in intermittent contact mode in air. MSNL-F (f = 120 kHz, k = 0.6 N m-1, tip radius of curvature < 12 nm) probes were used. The AFM topography images were leveled, line-corrected and measured (line profiles for diameter determination) using Gwyddion, a free and open-source SPM (scanning probe microscopy) data visualization and analysis program [41]. AFM samples were prepared on silicon substrates freshly cleaned in piranha solution. For bare and functionalized MSNPs that carry a negative surface charge, the clean substrate was first incubated in PAH, followed by rinsing and drying. On the other hand, PEI-modified MSNPs were deposited on a bare silicon substrate. For each sample, the nanoparticle suspension was incubated for 1 min on the substrate, before rinsing with ultrapure water and drying with pure nitrogen gas.

### Cell culture

A549 cells, HepG2, BJ1-hTERT, NIH3T3, Hek293 and A431 cells were cultured in 25 cm^2^ culture flasks at 37°C under 5% CO_2_ atmosphere. All cell lines were maintained in DMEM medium with 10% FBS, 1% L-glutamax and 0.1% gentamicin. For fluorescence microscopy experiments, the cells were seeded in 29 mm glass bottom dishes (Cellvis) and grown until ∼60% confluency before adding the nanoparticles.

### Immunofluorescence labelling

Cells were stained with both the dual-labelled and non-labelled CD44 antibody. First, cells were seeded in two 29 mm glass bottom dishes and grown overnight. Next, the cells were fixed with paraformaldehyde (4%) and the membrane was permeabilized with Triton X-100 (0.1%), for 10 minutes. The sample was carefully washed with PBS (1X) between each step. After washing, blocking was performed for 1 h with a bovine serum albumin solution (3% in PBS). The dual-labelled and non-labelled antibody were added to both the A549 and HepG2 cells at a final concentration of 2 μg/mL and incubated overnight at 4 °C. After antibody incubation, the dual-labelled antibody samples were washed 3 times with PBS. The sample containing non-labelled antibody were washed with 3 times with PBS and incubated with the secondary antibody, donkey-anti-rat IgG-AF488 a final concentration of 1 μg/mL for 2 h. After that, the samples were washed with PBS.

### Ab-PEI-MSNPs targeting efficiency

A549 cells, HepG2, BJ1-hTERT, NIH3T3, Hek293 and A431 cells were seeded in a 29mm glass bottom dish and grown until 60 – 80 % confluency. Fluorescent Ab-PEI-MSNPs were added to the cells to a final concentration of 50 µg/mL. After 6 h of incubation with the NPs, the cells were washed 3 times with PBS and fresh medium was added to the samples. The samples were placed back in the incubator for an additional 24 h incubation. Prior to imaging, the plasma membrane was stained using DiR (1 µM) in HBSS for 13 min and the sample was washed 3 times with HBSS.

### Fluorescence microscopy

Confocal fluorescence imaging was performed on a Leica TCS SP8 mini microscope implementing a HC PL APO 63x water immersion objective (NA 1.2). Distinct diode lasers were used depending on the dye. For LysoTracker™ DeepRed and DiR lipophilic dye, a red 638 nm diode laser was used at a laser power between 10 and 60 µW. For RhoB or Atto565 detection, a green 552 nm diode laser was used for excitation (laser power between 20 and 80 µW), while the blue 488 nm diode laser was used to excite Doxorubicin, FITC and Atto488 (laser power between 10 and 50 µW). The laser powers were measured at the objective. Detection was performed with HyD SMD high sensitivity detectors in standard mode, operating in a detection range of 400 to 800 nm. The detection range was adjusted depending on the dye, being 500-550 nm for Atto488 and FITC detection, 570-600 nm for Doxorubicin, RhoB and Atto565, and 650-750 nm for LysoTracker™ DeepRed and DiR lipophilic dye detection. The images were acquired with a z-step of 1 µm and line averaging of 3.

### αCD44-PEI-MSNPs intracellular localization

A549 cells were incubated with αCD44-PEI-MSNPs_RhoB for 24 h (at a final concentration of 50µg/mL). The sample was then washed 3 times with PBS to remove extracellular nanoparticles and kept in HBSS during image acquisition. One sample was imaged immediately after 24 h incubation, while two other replicates were placed back in the incubator for the 48 and 72 h time points (after replacing the PBS for cell culture medium). Prior to imaging, the samples were washed with HBSS and the lysosomes were stained with LysoTracker™ Deep Red (100 nM final concentration in HBSS) for 15 minutes. After washing 3 times with PBS, the samples were imaged in HBSS. The fluorescence images acquired (see *Fluorescence microscopy* section) were processed and analyzed using Fiji and the built-in co-localization plugin JACoP [42]. Within this plugin, Manders co-localization was found as an appropriate analysis strategy as this method measures the fraction of co-occurrence of the signal in two channels rather than their correlation [43]. After manual selection of the cell area with the region of interest (ROI) manager, the Manders’ coefficient (MC) was calculated for 28-40 biological replicates for each time point (using 3 technical replicates). More specifically, MC indicates the fraction of pixels of the αCD44-PEI-MSNPs ROI that overlap with the pixels of the LysoTracker ROI, resulting in a value between 0 and 1. One means that 100% of the pixels of the αCD44-PEI-MSNP ROI overlap with the pixels of the LysoTracker channel and 0 being a 0% pixel overlap.

### Doxorubicin release

αCD44-PEI-MSNPs were loaded with doxorubicin to a final concentration of 40 μM (αCD44-PEIMSNP_Dox). A549 cells were incubated with αCD44-PEI-MSNP_Dox for 24 h. After 24 h of incubation, the sample was washed with PBS 3 times to remove extracellular nanoparticles and kept in HBSS during image acquisition. One sample was immediately measured (with confocal fluorescence microscopy), while 2 other replicates were placed back in the incubator for the 48 and 72 h time points (after replacing the PBS by cell culture medium). An additional sample was checked after 8 h of nanoparticle incubation, in order to monitor nanoparticle endocytosis. Prior to imaging, the samples were washed 3 times with PBS and the acidic vesicles were visualized by adding a solution containing LysoTracker™ Deep Red (100 nM final concentration in HBSS) for 15 minutes. After a final washing step with PBS, the samples were imaged in HBSS using a confocal microscope.

### Anticancer efficiency

A549 cells were seeded in a 96-well plate at a density of 2 × 10^4^ cells/well. The next day, 25, 50, 100, 200 and nM of Dox and 25, 50, 100 and 200 μg/mL of the empty and Dox loaded αCD44-PEI-MNSPs were added to the A549 cells. Four biological replicates were prepared for each condition. Cells incubated with nanoparticles were washed with PBS after 24 h of nanoparticle incubation to remove the excess of nanoparticles. The PBS was then replaced by fresh medium and the sample was placed in the cell incubator. 72 h after the addition of free Dox or nanoparticles, the cells were washed 3x with PBS and fixed with 4% PFA (in PBS). Cells were incubated with Hoechst 33342 (1μg/mL) for 1 h. After washing with PBS, the viable cells were imaged using a Lionheart FX automated microscope (BioTek) implementing a 10x air objective (NA: 0,3), a high power LED of 365nm, combined with a DAPI filter cube. Images were analyzed using the Gen5™ software. *Statistical analysis*. The data are displayed as means ± standard deviations and error bars indicate ± standard deviation. A randomization test was used to compare any two groups of values and performed in the online software tool “Plots of Difference” [44]. Statistical significance was reported as * p < 0,05, ** p < 0,01, *** p < 0,001

## 3. Results and Discussion

### 3.1. αCD44 conjugated nanoparticles

CD44 is a transmembrane glycoprotein receptor overexpressed in different cancers (e.g. breast, lung, colon, head and neck and pancreatic cancer [45–48]) and cancer stem cells [49,50]. Its presence is often associated with high malignancy and chemo-resistance, making it an important cancer biomarker and target for cancer therapy. Hyaluronic acid (HA) is a ligand for the CD44 receptor, and has therefore been widely exploited as a polymer coating in targeted DDSs[2,51,52]. The ligand, HA, is naturally present in the extracellular matrix (ECM), leading to possible competition between HA from the ECM and the HA on the nanoparticles. As an alternative, antibodies with a high affinity for CD44 can be implemented in the DDS to target the receptor [53–55].

#### 3.1.1. Synthesis of Azide-functionalized MSNPs

Our approach to conjugate antibodies to nanoparticles comprise the grafting of different groups on both the antibody and the nanoparticle, separately (Figure 1). As a model for nano-carriers, we used MSNPs. PEI-coated MSNPs were prepared as previously described [56]. Briefly, MSNPs were synthesized using the biphase stratification method and loaded with either Dox or RhoB, for cytotoxicity studies or imaging assays, respectively. The fluorescence spectra of the dye/drug loaded MSNPs are shown in Figure S1. A PEI layer (Mw = 1.3 kDa) was deposited on the MSNPs (PEI-MSNPs), through electrostatic interactions. In our approach, the PEI layer serves two purposes: to provide the nanoparticles with the capability of endosomal escape (as reported by *Fortuni et al*. [56]) and to functionalize the surface of the NPs with amine groups. These amine groups were used as an anchor for further covalent functionalization. An azide moiety was covalently linked to the PEI-MSNPs making use of an NHS ester-PEG_4_-N_3_ linker via NHS ester coupling (N_3_-PEI-MSNPs, Figure 1a). The central poly(ethylene)glycol (PEG) chain provided extra stealth to the DDS, increasing its biocompatibility. The resulting surface azide groups served as docking sites for the antibodies.

#### 3.1.2. Labelling of CD44 antibodies

To attach the antibody molecules to the azide-grafted nanoparticles, the antibodies were functionalized with a DBCO moiety (Figure 1b). Additionally, for visualization purposes, a fluorescent dye (Atto488) was also added, yielding dual-labelled antibodies. As described in the Methods section, labelling of the lysine residues of the antibody was achieved via an NHS ester coupling reaction. To evaluate if the addition of DBCO and/or Atto488 affect the specificity of the antibody, we performed immunostainings of two cell lines, A549 and HepG2 cells, with high and low expression level of CD44, respectively [45][57]. Before testing the specificity of the DBCO/Atto488 labelled antibodies, the CD44 expression level in both cell lines was evaluated using immunofluorescence. For this, we stained the cells with unmodified CD44 antibody, followed by a second staining step with a fluorescently labelled secondary antibody. As shown in Figure 2a, CD44 receptor molecules were detected in A549 cells (localized to the cell membrane), while no CD44 immunostaining was visible on the fluorescence images acquired for HepG2 cells. A549 cells stained with the DBCO/Atto488 dual labelled CD44 antibodies displayed a fluorescence signal on the plasma membrane, indicating that the specificity of the antibody was retained after labelling (Figure 2b). The fluorescence signal present in the nucleus was attributed to the free dye molecules still present in solution after the labelling procedure (Figure 2c).

**Figure 2:**
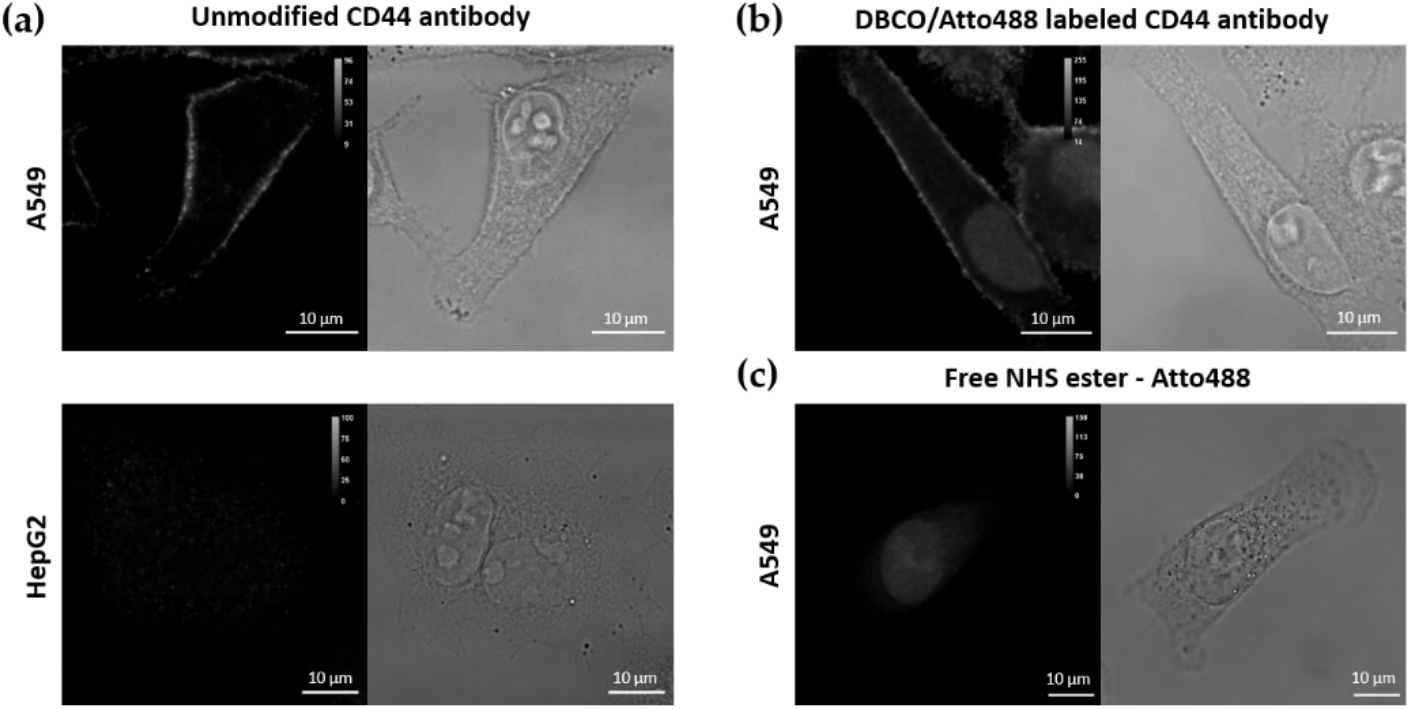
Immunofluorescence (IF) using (a) the unmodified primary CD44 antibody and a secondary labelled antibody on A549 and HepG2 cells, (b) dual DBCO/Atto488 labelled CD44 antibodies in A549 cells and (c) free NHS ester-Atto488 molecules. Scale bar is 10 µm.

#### 3.1.3. Synthesis and characterization of αCD44 conjugated MSNPs

The antibody was covalently linked on the nanoparticle via copper-free click chemistry between the azide moiety and the DBCO-labelled-antibody (generating αCD44-PEI-MSNPs, Figure 1c). The developed nanoparticles were characterized using scanning electron microscopy (SEM), atomic force microscopy (AFM), zeta potential measurements and fluorescence microscopy.

SEM images displayed a uniform size and shape homogeneity of the bare MSNPs, PEI-MSNPs and αCD44-PEI-MSNPs (conjugation with αCD44), with a mean diameter of 118, 119 and 124 nm, respectively (Figure 3a). The average size of the bare and PEI-coated MSNPs is in agreement with previous reports [56]. Further, a low degree of aggregation was observed in all the samples. Although there was no significant difference in diameter between the bare MSNPs and the PEI-MSNPs, a significant increase could be observed upon antibody functionalization (p < 0,05). These results were confirmed with AFM measurements, where an average nanoparticle height of 121, 123 and 128 nm was detected for the MSNPs, PEI-MSNPs and αCD44-PEI-MSNPs, respectively. The corresponding AFM images and plots are displayed in supporting information (Figure S2). The increase in MSNP height upon antibody functionalization indicated a successful attachment to the surface. The small difference can be attributed to the average size of an antibody (in the range of 5 to 15 nm depending on its orientation).

**Figure 3:**
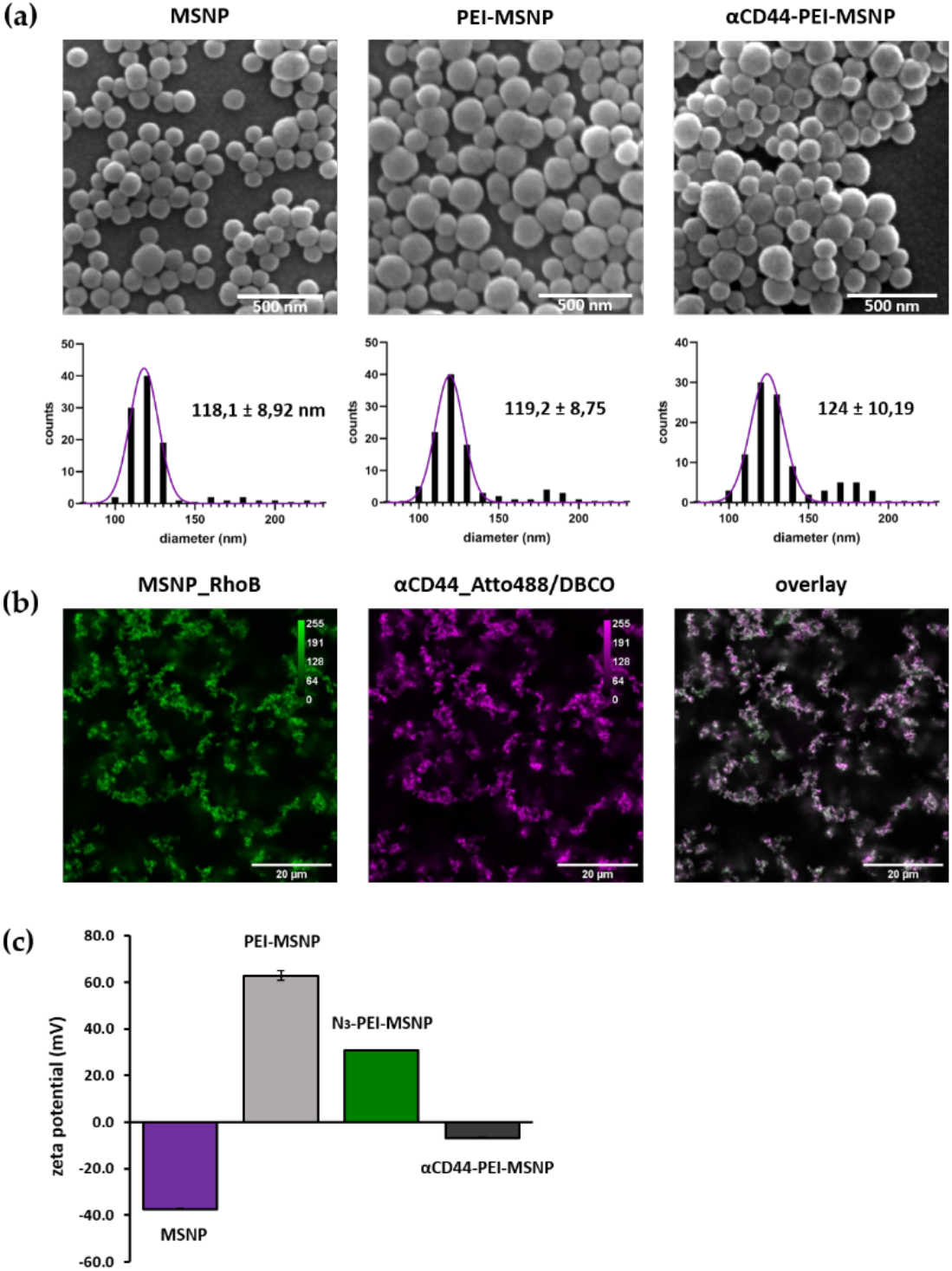
Characterization of bare (MSNPs), PEI-coated (PEI-MSNPs), Azide-functionalized (N_3_-PEI-MSNPs) and CD44 conjugated (αCD44-PEI-MSNPs) mesoporous silica nanoparticles. (a) Representative SEM images and size distribution of the imaged particles. Values shown as mean ± SD). Scale bar is 500 nm. (b) Confocal fluorescence images of αCD44-PEI-MSNPs in which MSNPs were loaded with RhoB (first panel, green), while an Atto488 (and DBCO) label was conjugated to the CD44 antibody (second panel, magenta). An overlay is displayed in the third panel. Scale bar is 20µm (c) Zeta-potential measurements given as mean ± SD.

Zeta potential measurements were used to follow each step of the functionalization process. Bare MSNPs had a negative charge (−37,5 mV) due to the partially deprotonated hydroxyl groups on the MSNP surface. Upon PEI coating, the zeta potential increased to +63 mV as a result of the presence of amine groups in PEI. Subsequent coupling of the azide moiety was reflected in a decrease in the zeta potential (+31 mV) (Figure 3c). This decrease can be associated to the negative charge of the N_3_ group, which partly compensates the charge from amino groups of PEI. Finally, a further decrease in zeta potential (−6,7 mV) upon antibody grafting could be observed (Figure 3c). The charge of the nanoparticle after antibody conjugation can be related to the isoelectric point (IEP) of the antibody. Since the IEP of IgG antibodies lies between 6,6 and 7,2 [58], antibodies are expected to be negatively charged at neutral pH, which agrees with our zeta potential results and previously reported results [31,59]. To evaluate the uniformity of antibody conjugation, MSNPs were loaded with a dye (RhoB) before functionalization. Since the antibodies were fluorescently labelled (Atto488), the colocalization between the RhoB-loaded MSNPs and the antibody was evaluated with confocal fluorescence microscopy. Figure 3b shows a clear overlap between the atto488-labelled antibody and the RhoB loaded MSNPs. This fluorescence data supports the findings obtained from AFM and the zeta potential measurements, indicating the proper conjugation of the antibody.

### 3.2. Selectivity and efficiency of αCD44 functionalized particles

#### 3.2.1. Targeting capability of αCD44-PEI-MSNPs

To assess the targeting capability and specificity of the antibody conjugated MSNPs towards CD44-overexpressing cells, RhoB-loaded MSNPs were used to monitor the cellular uptake via fluorescence microscopy. The targeting efficiency was tested by comparing the number of nanoparticles internalized in A549 cells (human lung carcinoma cells, with CD44 receptor overexpression [60]) and HepG2 cells (liver carcinoma, with low CD44 receptor expression [61]). Both cell lines were incubated for 6 h with MSNPs presenting different functionalizations: no coating (MSNPs), only a PEI coating (PEI-MSNP) and both the PEI and antibody functionalization (αCD44-PEI-MSNPs). Afterwards, the medium was refreshed to avoid further nanoparticle internalization and the cells were incubated overnight (24 h incubation in total). The internalization was quantified by confocal fluorescence imaging after staining the plasma membrane with a lipid intercalating dye (DiR). The fluorescence images are shown in Figure 4. The uptake of bare MSNPs was minimal in both cell lines (Figure 4a and 4d), while a coating with PEI resulted in an increase in internalization in both A549 and HepG2 (Figure 4b and 4e). The increase in cellular uptake of nanoparticles upon PEI functionalization is in agreement with previous reports and can be attributed to the positive charge of PEI-MSNPs, leading to a higher interaction with the negatively charged plasma membrane, which facilitates nanoparticle internalization [56,62–64]. It is important to note that this enhanced uptake of PEI-coated MSNPs was cell line unspecific. Thanks to the charge drop (from +63 mV to -6,7 mV), this unspecific internalization was minimized upon antibody conjugation to MSNPs. As such, uptake of αCD44-PEI-MSNPs in HepG2 was lower, similar to the internalization of bare MSNP (Figure 4f). On the other hand, A549 cells incubated with αCD44-PEI-MSNPs displayed a high number of internalized particles, especially when compared to bare MSNP (Figure 4c). A quantitative analysis on the differential internalization of αCD44-PEI-MSNPs in A549 and HepG2 cell was performed resulting in a significant difference in normalized mean intensity (being 0,31 and 0,017 for A549 and HepG2 cells, respectively) (see Methods and Figure S3).

**Figure 4:**
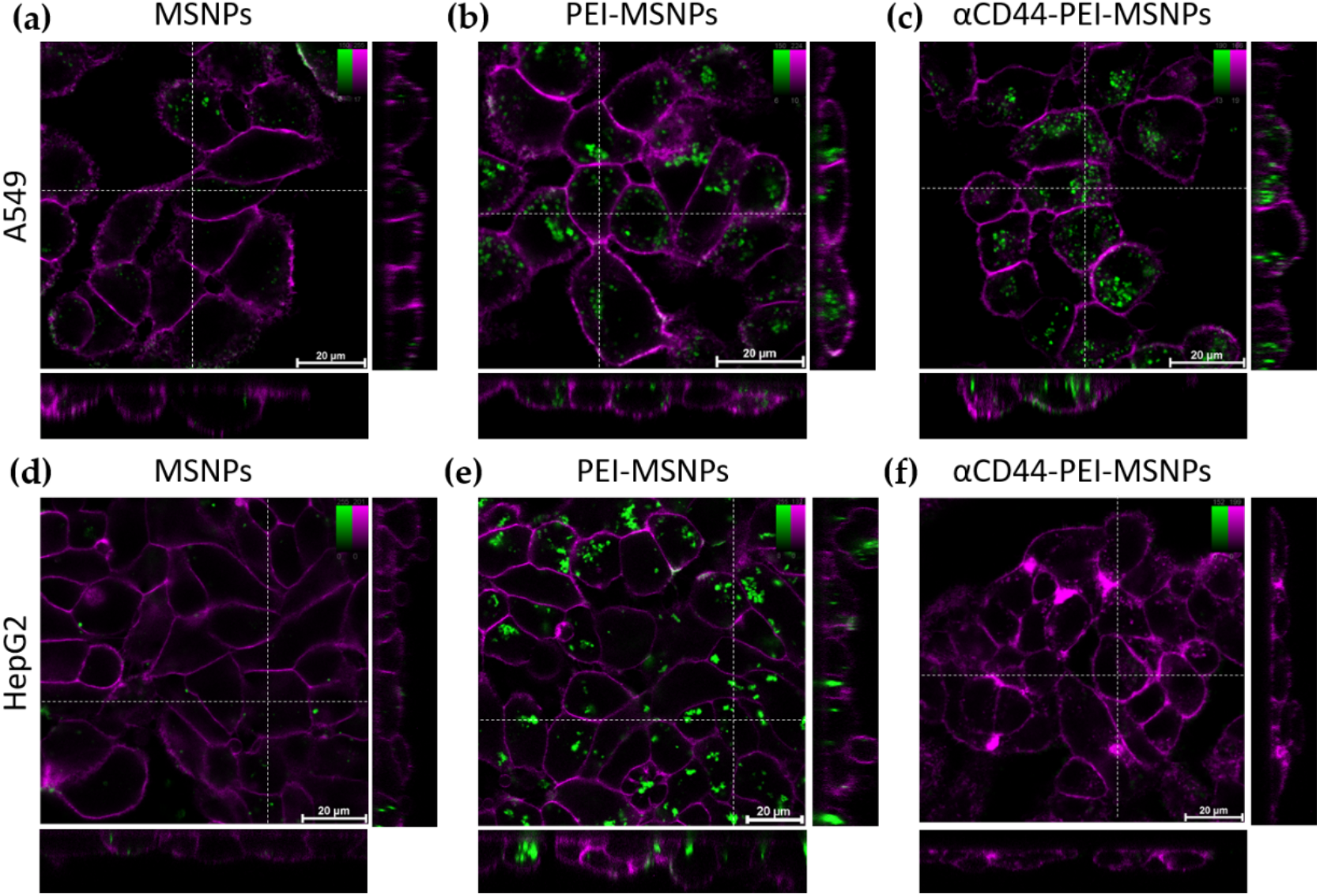
Confocal fluorescence microscopy images showing the influence of different MSNP coatings on the uptake of nanoparticles in A549 and HepG2 cells. Internalization of bare MSNP (a,d), PEI-coated MSNPs (PEI-MSNP, panels b,e) and CD44 functionalized MSNPs (αCD44-PEI-MSNP, panels c,f) in A549 cells (a-c) and HepG2 cells (d-f). Nanoparticles were loaded with RhoB (green) and the plasma membrane was stained with DiR (magenta). The central square represents a single xy plane, while the bottom and left panels are the xz and yz cross-sections, indicated by the dashed lines. Scale bar is 20 μM, color bars display the intensity values.

To assure that the observed increase in cellular uptake in CD44-overexpressing cells was not linked to the specific cell lines used, we checked the uptake of αCD44-PEI-MSNPs in two fibroblast cell lines, NIH 3T3 (mouse embryonic fibroblasts with a low CD44 expression [65,66]) and BJ1-hTERT (human fibroblasts with high CD44 expression [67,68]), using the same experimental approach (Figure S4). The respective CD44 expression in the two cell lines was validated with a standard immunofluorescence staining of CD44 (Figure S5). The discrepancy between the uptake of αCD44-PEI-MSNPs in fibroblasts with different expression levels of CD44 confirmed the targeting efficiency of our nanoparticles after conjugation with the CD44 antibody.

#### 3.2.2. Intracellular trafficking

One of the main bottlenecks faced by DDSs is their entrapment in acidic vesicles and subsequent degradation, significantly limiting the overall efficiency of the DDS [69–71]. To this end, strategies have been developed to facilitate an endosomal escape, releasing the nanoparticles (or their cargo) into the cytoplasm. We have previously shown that the addition of a PEI shell leads to the release of the nanoparticles in the cytoplasm [56]. To verify that conjugation to an antibody does not affect the endosomal escape capability of the PEI layer, the co-localization of αCD44-PEI-MSNPs with lysosomes was monitored using fluorescence microscopy. Briefly, A549 cells were incubated with RhoB-loaded αCD44-PEI-MSNPs for 24 h, after which the medium of the samples was refreshed, in order to avoid further nanoparticle internalization (i.e. only nanoparticles that were endocytosed within the first 24 h of incubation were followed). At each time point (24, 48 and 72 h after particle addition), the acidic vesicles were stained with LysoTracker Deep Red and the samples were imaged. Figure 5 shows representative fluorescence images at the different time points. To quantify the co-localization between the acidic vesicles and the αCD44-PEI-MSNPs through time, the Manders’ co-localization coefficient was calculated (Figure <u>S6</u>). Briefly, an intensity-based threshold was used to calculate the areas of the image corresponding to nanoparticles and to lysosomes. The Manders’ coefficient (MC) calculates the degree of overlap between objects in different channels (with 0 being no overlap, and 1 indicating a complete overlap). Fluorescence images showed that, within 24 h, almost all nanoparticles were located inside the acidic vesicles (Figure 5a). While at 24 and 48 h, no relevant escape was observed with MC of 0,88 and 0,82 (Figure S6), a significant decrease in the co-localization of nanoparticles and lysosomes was found after 72 h (MC of 0,66), indicating that after 72 h a considerable number of αCD44-PEI-MSNPs were localized outside the lysosomes (Figure 5c). Overall, these results suggest that the presence of αCD44 on the surface of the nanoparticles does not affect the endosomolitic activity of the PEI coating as most of nanoparticles were able to dissociate from the lysosomes within 72 h. This was validated by the significant difference in MC found between 48 and 72 h (Figure S6).

**Figure 5:**
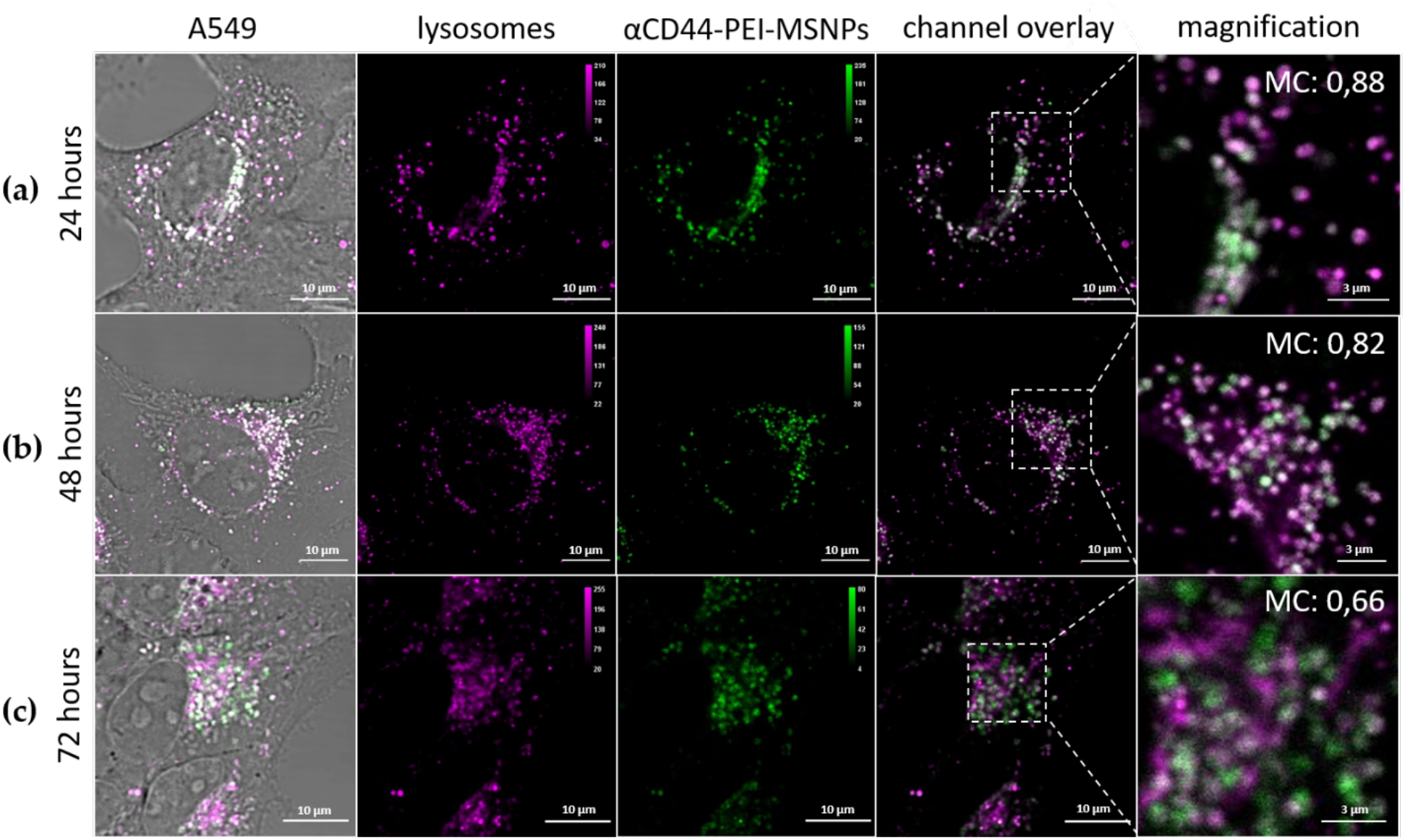
Intracellular localization of RhoB-loaded αCD44-PEI-MSNPs (green) with respect to the lysosomes (Lysotracker Deep Red, magenta) over time. A549 cells are incubated with αCD44-PEI-MSNPs for (a) 24 h (b) 48 h and (c) 72 h. The first column shows a complete merge including the transmission image, also displaying the cell areas. In the second and third column, the lysosomes and nanoparticles are depicted, in purple and green respectively. In the fourth and fifth column, a channel overlay and respective magnification are shown. Scale bar is 10 μm in main images and 3 μm in the magnified images.

#### 3.2.3. Intracellular release of doxorubicin

As shown in the previous section, a considerable number of the internalized αCD44-PEI-MSNPs were able to escape the acidic vesicles. Aside from the ability to escape the lysosomes, a higher efficiency requires that enough cargo can be released into the cytoplasm in a controlled fashion. To monitor the drug release after cellular uptake, doxorubicin was used as drug model and encapsulated in the pores of MSNPs prior to coating (αCD44-PEI-MSNPs_Dox). Dox is a cytostatic anticancer drug that is used to treat different types of cancer, for instance leukemia, lymphoma and breast cancer. Its mechanism of action is based on the intercalation with the DNA, resulting in cell death [72]. Due to the fluorescent nature of Dox, its intracellular localization could be monitored over time with confocal fluorescence microscopy. To show that αCD44-PEI-MSNPs_Dox carried Dox (cyan, Figure 6) into the target cell and exhibited a controlled intracellular drug release, A549 cells were incubated with αCD44-PEI-MSNPs_Dox for 8, 24, 48 and 72 h. To avoid further nanoparticle internalization, the medium of the samples was refreshed 24 h after the addition of the nanoparticles. The acidic compartments were stained with LysoTracker Deep Red (magenta, Figure 6) to analyze the cargo (Dox, cyan) release/leakage from the lysosomes. While after 8 and 24 h, most of the Dox was still retained inside lysosomes (indicated by the high overlap between the Dox and the lysosomes) (Figure 6a-b), increasing amounts of Dox discharge into the cytosol were observed after 48 and 72 h (Figure 6c-d). The overlap between the acidic vesicles and Dox molecules at 8 and 24 h indicated that at this time point, Dox was presumably still inside the pores of the MSNPs or being slowly released inside the acidic vesicles. At an acidic pH, the silica hydroxyl groups are protonated, hindering the electrostatic interaction between the PEI and the silica surface. Therefore, we hypothesize that the decreased pH inside the lysosomes aids in the release of the shell, and facilitates the consequent Dox release from the mesoporous silica pores. Accordingly, increasing amounts of Dox were detected in the cytoplasm over 48 to 72 h (Figure 6c-d). Moreover, as shown in Figure 6d, the cells morphology suggested cellular death, indicating that the cells were being killed by the successful release of the drug.

**Figure 6:**
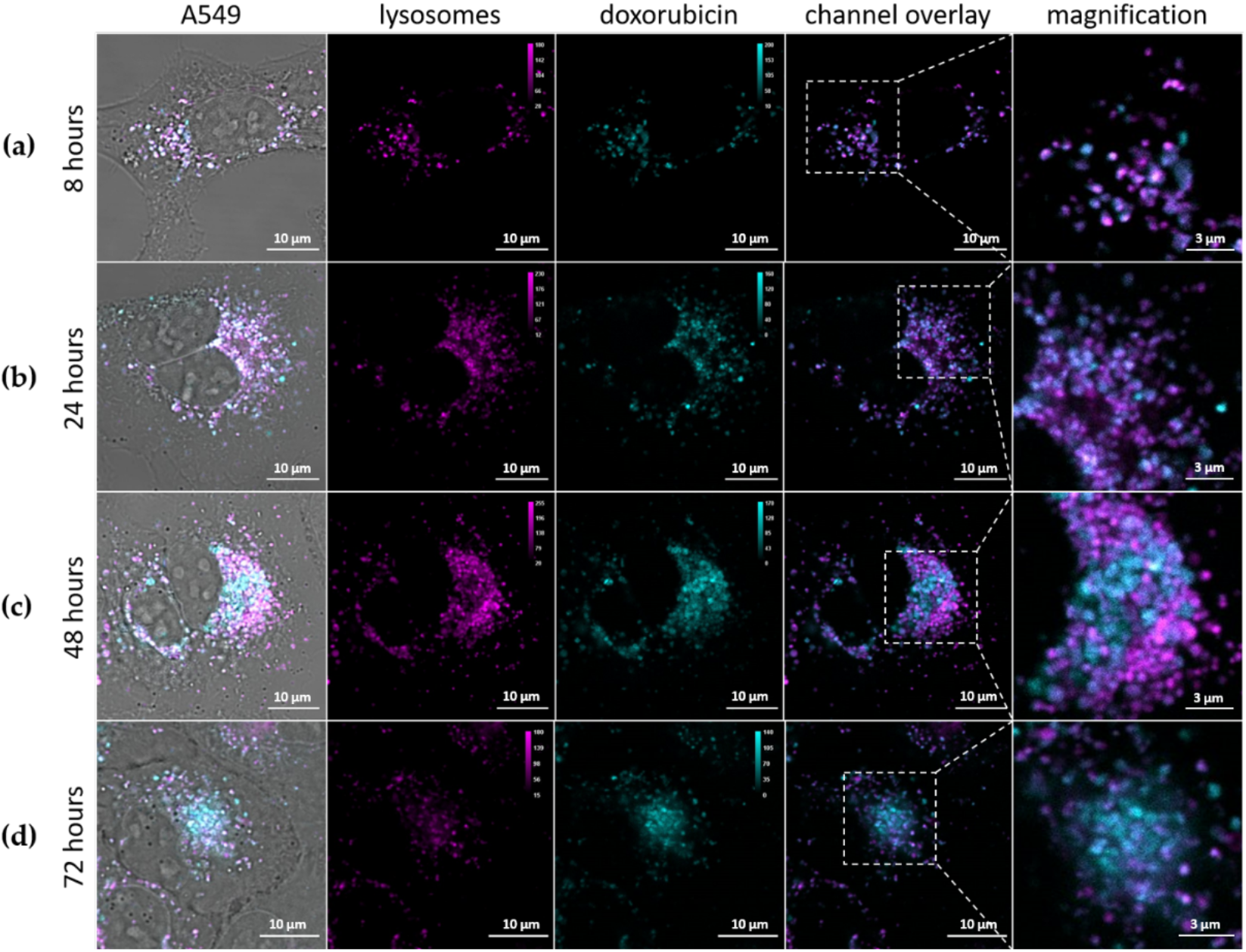
Intracellular release of Dox (cyan) from αCD44-PEI-MSNPs_Dox with respect to the lysosomes (magenta) over time. A549 cells are incubated with αCD44-PEI-MSNPs_Dox for (a) 8 h (b) 24 h (c) 48 h and (d) 72 h. The first column shows a complete merge including the transmission image, also displaying the cell contours. In the second and third column, the lysosomes and Dox are depicted, in pink and cyan, respectively. In the fourth and fifth column, a channel overlay and respective magnification are shown. Scale bar is 10 μm in main images and 3 μm in the magnified images.

#### 3.2.4. Viability studies

To evaluate the toxicity and anti-cancer efficiency of the antibody coated MSNPs, the viability of A549 cells was investigated after 72 h of incubation with pure Dox, Dox-loaded nanoparticles (αCD44-PEI-MSNPs_Dox) and empty nanoparticles (αCD44-PEI-MSNPs). After 24 h of incubation with the nanoparticle/drug solution, the cells were washed to remove excess nanoparticles. Different concentrations of Dox and nanoparticles were added ranging from 25 nM to 200 nM for free Dox and 25 to 100 μg/mL for the nanoparticles (Figure 7). While empty αCD44-PEI-MSNPs could be considered non-toxic, reaching a minimum of 92% cell viability at the highest concentration (100 µg/mL), Dox loaded αCD44-PEI-MSNPs exhibited cytotoxic effects, reaching a minimum of 38% viability at the same concentration. The cell viability in the presence of the drug loaded nanoparticles decreased with increasing nanoparticle concentration while the viability with the empty nanoparticles stayed roughly constant. Similar to the drug loaded nanoparticles, a dose dependent response on the cell viability was observed for free Dox. However, we found an evident difference in the cell killing effect between the free Dox and the Dox loaded nanoparticles (p < 0,05). This enhanced cytotoxicity is potentially due to a higher internalization rate of the αCD44-PEI-MSNPs followed by an endosomal release and specific drug release in the cytoplasm.

**Figure 7:**
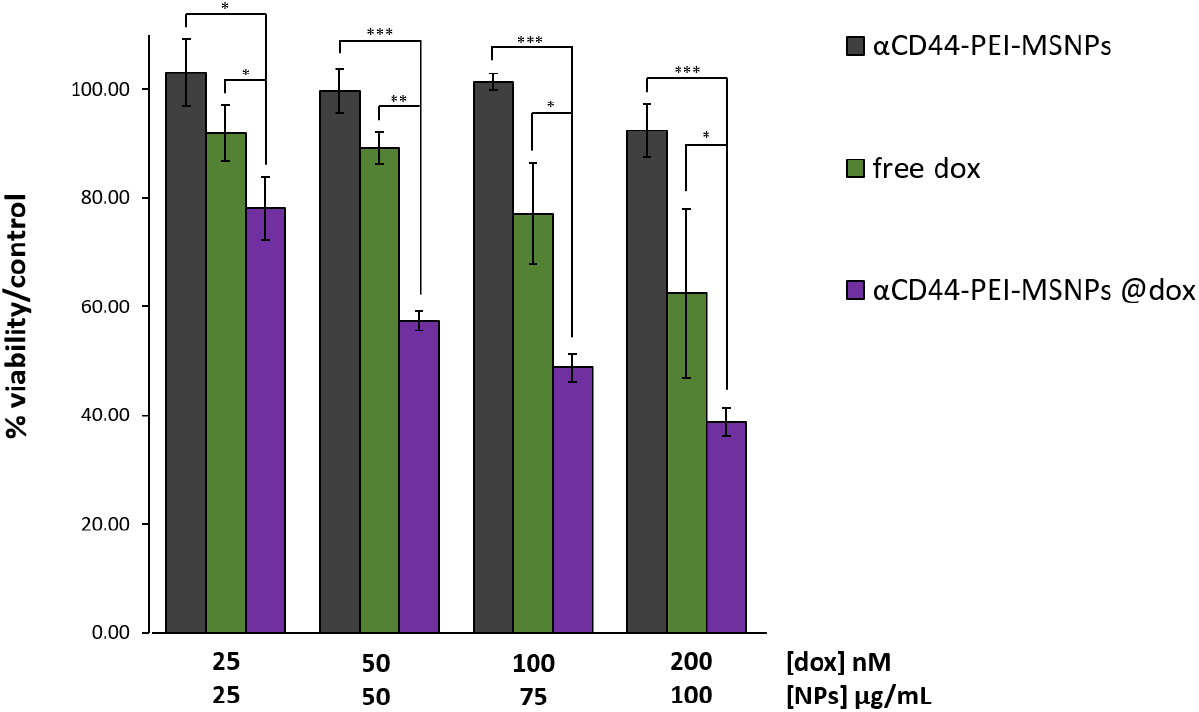
Viability of A549 cells after 72h incubation with different concentrations of free Dox, Dox loaded αCD44-PEI-MSNPs and empty αCD44-PEI-MSNPs. Dox concentrations are expressed in nM while nanoparticle concentrations are in µg/mL. Error bars indicate ± SD, with *(p < 0,05), **(p < 0,01) and ***(p < 0,001), n=4.

From these results, it can be concluded that Dox loaded αCD44-PEI-MSNPs exhibited high toxicity to CD44 overexpressing cells (higher than free Dox), while only limited toxicity was observed for the empty carrier. Consequently, a satisfying therapeutic efficiency can be concluded, through an efficient nanoparticle internalization and high drug payload delivery targeted to the cancer cell cytosol.

### 3.3. αEGFR conjugated nanoparticles

To prove the versatility of our antibody coating strategy, we targeted next a second cancer marker, EGFR. EGFR overexpression is related to different types of cancers and is often associated to a poor prognosis for the patient. Especially in glioblastoma, lung and breast cancer, where EGFR stimulates tumour growth [34]. To this end, EGFR has emerged as a popular target for cancer cell specific therapies. As a result, many nanoparticles targeting EGFR were developed, either via nanoparticle conjugation with its ligand, EGF, or by attaching EGFR antibodies [73–76].

#### 3.3.1. Labelling of the EGFR antibodies

For labelling the EGFR antibodies, the same method was used as for the CD44 antibodies (section 3.1.2). Similarly, labelling with DBCO was performed, enabling the conjugation of DBCO-EGFR antibodies to the azide-coated nanoparticles. For fluorescence microscopy purposes, the EGFR antibodies were also conjugated to a dye (Atto565). The dual-labelling of the EGFR antibodies with DBCO and Atto565 was carried out via an NHS ester coupling reaction with the antibodies’ lysine residues (as described in Methods). We checked whether the specificity of the EGFR antibody was retained after dual-labelling (with Atto565 and DBCO) by performing immunofluorescence on two cell lines, A431 and Hek293 cells, presenting high and low expression of EGFR, respectively [77,78]. First, the EGFR expression in the chosen cell lines, A431 and Hek293 was determined via immunofluorescence with the unmodified EGFR antibody (followed by staining with a fluorescently labelled secondary antibody). As shown in Figure 8a, EGFR was visualized on the plasma membrane of A431 cells, while no EGFR could be detected in Hek293 cells. A431 cells stained with the DBCO/Atto565 labelled EGFR antibody also displayed the plasma membrane-localized signal, proving that the specificity of the EGFR antibody was not hindered by our labelling protocol (Figure 8b). A nuclear background could be observed, associated to free Atto565 molecules, similar to what was observed for the Atto488 molecules (see section 3.1.2.).

**Figure 8:**
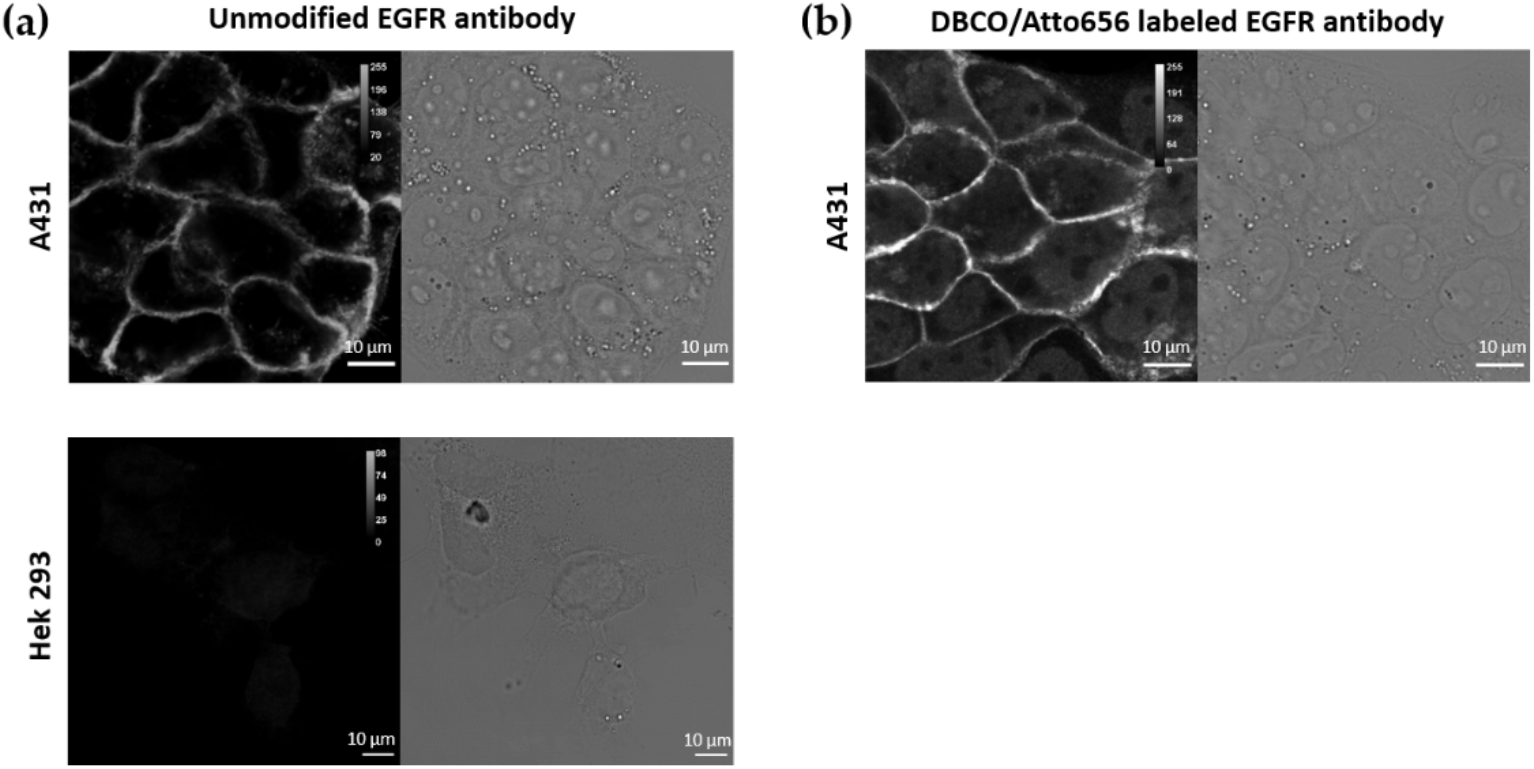
Immunofluorescence (IF) using (a) the unmodified primary EGFR antibody followed by a labelled secondary antibody on A431 and Hek293 cells, (b) DBCO/Atto565 labelled EGFR antibodies in A431 cells. Scale bar is 10 µm.

#### 3.3.2. Synthesis and characterization of αEGFR conjugated MSNPs

The EGFR antibody was attached to the Azide coated nanoparticles via the same approach (αEGFR-PEI-MSNPs). Since the EGFR antibody is labelled with an Atto565 dye (as described in the previous section), to avoid spectral overlap during fluorescence microscopy studies, azide conjugated nanoparticles were labelled with Fluorescein (FITC). FITC was covalently attached to the silica matrix, as stated in the Methods section. The resulting αEGFR conjugated nanoparticles were characterized via SEM, AFM, zeta potential measurements and fluorescence microscopy.

Similar to αCD44-conjugates particles, αEGFR-PEI-MSNPs displayed a uniform size and shape homogeneity (Figure 9a). With a mean diameter of 122 nm (as derived from the SEM data), αEGFR-PEI-MSNPs were significantly bigger than the PEI-MSNPs (Figure 3a) (p < 0,05), because of the EGFR layer. AFM measurements revealed an average nanoparticle diameter of 127 nm, which agreed with the SEM results (Figure S3). Zeta potential measurements were performed to check the different steps of nanoparticle functionalization. The zeta potential data of the bare MSNPs, PEI grafted MSNPs (PEI-MSNPs) and Azide conjugated MSNPs (N_3_-PEI-MSNPs) were already discussed in section 3.1.3 (Figure 3c). αEGFR-PEI-MSNPs displayed a negative zeta potential (−11,6 mV), similar to the one of the MNSPs after conjugation with the CD44 antibody (αCD44-PEI-MSNPs, -6,7 mV) (Figure 9c). Additionally, in order to confirm uniform binding of the EGFR antibody, MSNPs were encapsulated with a fluorescent dye (FITC), while the EGFR antibody was labelled with another dye (Atto565). Accordingly, the co-localization between the EGFR antibody and the MSNPs could be determined via confocal fluorescence microscopy measurements. This data confirms proper attachment of the EGFR antibody and uniform coverage of the nanoparticles with antibody molecules (Figure 9b).

**Figure 9:**
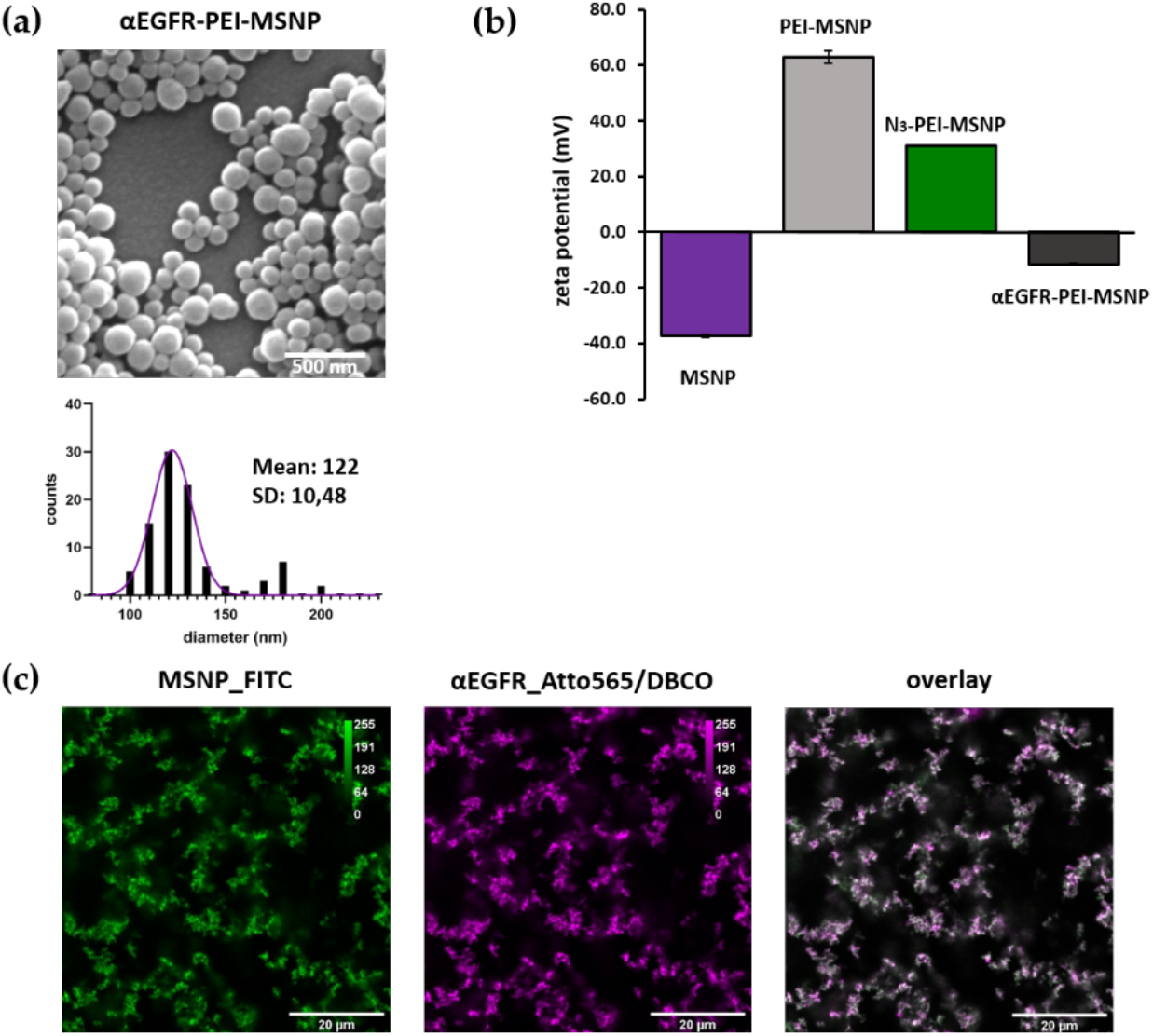
Characterization of EGFR conjugated (αEGFR-PEI-MSNPs) mesoporous silica nanoparticles. (a) Representative SEM image and size distribution of the imaged particles (values shown as mean ±SD). Scale bar is 500 nm. (b) Zeta-potential measurements. (c) Confocal fluorescence images of sedimented nanoparticles on glass showing the overlay (third panel) of FITC encapsulated MSNPs (first panel, green) and the Atto565/DBCO labelled EGFR antibody (second panel, magenta).

#### 3.3.3. Targeting capability of αEGFR-PEI-MSNPs

The uptake of nanoparticles with different coatings (MSNPs, PEI-MSNPs and αEGFR-PEI-MSNPs) was compared between A431 (epidermoid carcinoma with high EGFR expression [73]) and Hek293 cells (human embryonic kidney cells with low EGFR expression [78]). In agreement with the results of the preceding experiment, bare MSNPs were internalized in A431 and Hek293 in low amount (Figure 10a and 10d) and a similar increase in uptake could be detected for MSNPs with a PEI layer (Figure 10b and 10e). As predicted, conjugation of the EGFR antibody resulted in a discrepancy in particle uptake between A431 and Hek293 cells. Hek293 displayed a similar uptake behavior for the αEGFR-PEI-MSNPs and the bare MSNPs (Figure 10f) while in A431, there was an obvious increase in the number of αEGFR-PEI-MSNPs internalized, especially when compared to bare MSNPs (Figure 10c). This increase can be attributed to the specific recognition of the EGFR receptor by the αEGFR-PEI-MSNPs, proving cell specificity of these nanoparticles.

**Figure 10:**
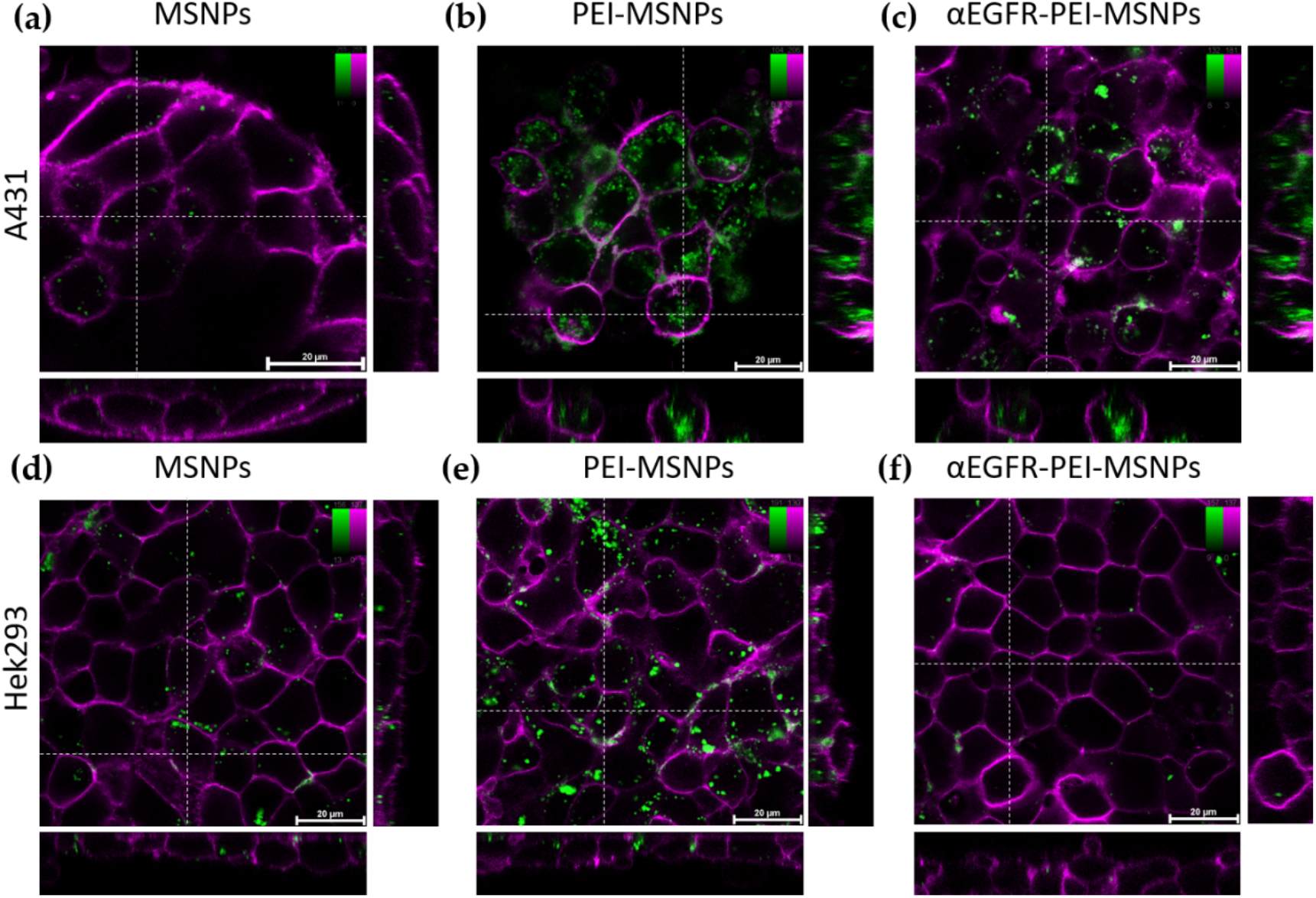
Confocal fluorescence microscopy stacks showing the influence of different MSNP coating in A431 and Hek293 cells. (a-c) Internalization of MSNP, PEI-MSNP and αEGFR-PEI-MSNP in A431 cells (from left to right). (d-f) Internalization of MSNP, PEI-MSNP and αEGFR-PEI-MSNP in Hek293 cells (from left to right). Nanoparticles (encapsulated with fluorescein) are displayed in green and the plasma membrane is stained with DiR (magenta). Scale bar is 20 μM, color bars display the intensity values.

## 5. Conclusions

In this work we show a facile method for the conjugation of different antibodies to nanoparticles using copper free click chemistry. Here, mesoporous silica nanoparticles (MSNPs) were chosen as drug carriers and the base for further functionalizations, however other nanoparticles can be used. Similarly, the Doxorubicin loaded inside the MSNPs can be substituted by other drugs for different therapeutic applications.

The first step in our approach was to coat the nanoparticles with a PEI layer. The amine groups provide the anchor for the covalent attachment of the azide moiety, to which the DBCO labelled antibodies were covalently linked via a simple click reaction. Importantly, click chemistry can be carried out under physiological conditions, without using any catalyst. Furthermore, the presence of the PEI layer reduced the effect of nanoparticle entrapment in the acidic vesicles (by taking advantage of the proton sponge effect) and supported the controlled drug release into the cancer cell.

For the conjugation to the nanoparticles, the antibodies were functionalized with a DBCO moiety. Since the antibodies were mono labelled with DBCO, our method prevents antibody polymerization and thus nanoparticle aggregation. Moreover, as this conjugation approach only requires the presence of amine groups at the surface of the antibody, the antibody itself can easily be adapted (based upon the expressed cancer marker). Antibody-conjugated DDS have therefore the potential to be extremely versatile.

The versatility of our coating strategy was demonstrated with two different antibodies, a CD44 -and an EGFR antibody, both showing excellent selectivity towards CD44 and EGFR overexpressing cells, respectively. This simple method can significantly contribute to the field of personalized cancer therapy, where the treatment is customized according to the cancer markers present in the tumour. In this regard, a variety of antibodies can be easily “clicked-on” for efficient targeting purposes.

## 6. Patents

### Supplementary Materials

The following are available online at www.mdpi.com/xxx/s1, Figure S1 to S6

## Author Contributions

Conceptualization, I.V.Z, B.F. and S.R.; methodology, I.V.Z., M.B., P.C., O.D. and B.F.; formal analysis, I.V.Z. and O.D.; investigation, I.V.Z., M.B., P.C., O.D., B.F. and S.R.; writing—original draft preparation, I.V.Z., M.B. and S.R.; writing—review and editing, all authors.; supervision, C.B., H.U., J.H., B.F. and S.R.; funding acquisition, C.B., H.U., J.H. and S.R. All authors have read and agreed to the published version of the manuscript.

## Funding

This work was supported by the Research Foundation—Flanders (FWO, projects G0A5817N, G0D4519N, 1529418N and G094717N), by KU Leuven (C14/16/053, C14/18/061 and KA/20/026). This collaborative work was also financially supported by the JSPS “Core–to–Core Program A. Advanced Research Networks”. I.V.Z. and B.F. acknowledge the support from FWO for their PhD and Postdoctoral fellowships, respectively (11F5419N and 12×1419N). P.C., J.H., H.U. and S.R. acknowledge financial support from the SuperCol project that received funding from the European Union’s Horizon 2020 research and innovation program under the Marie Skłodowska-Curie grant agreement No. 860914. J.H. further acknowledges the Flemish government through long-term structural funding Methusalem (CASAS2, Meth/15/04).

## Data Availability Statement

The data presented in this study is available on request from the corresponding authors.

## Acknowledgments

The authors acknowledge Dr. Roel Hammink (Radboud University, Nijmegen, NL) for the fruitful discussions concerning antibody functionalization. Figure 1 created with Biorender.com

## Conflicts of Interest

There are no conflicts of interest regarding this research.

## Notes

### Competing Interest Statement

The authors have declared no competing interest.

