## Supplementary info for "Versatile and Robust method for Antibody Conjugation to Nanoparticles with High Targeting Efficiency"

### Appendix A

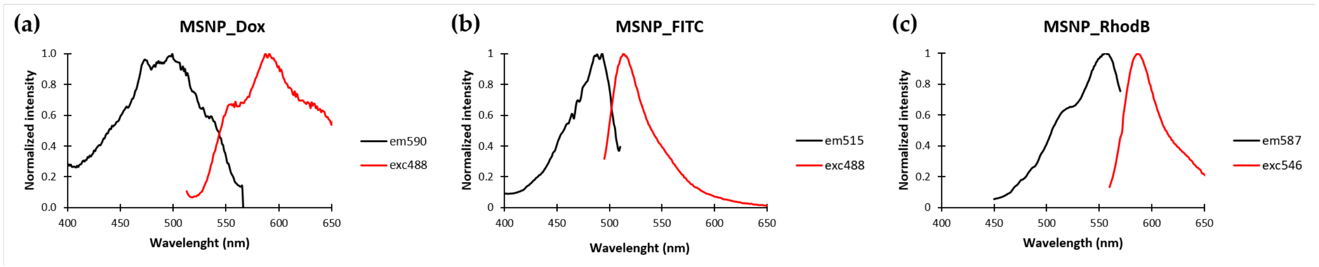


**Figure S1:** Emission and excitation spectra of drug and dye-loaded MSNPs. (a) dox-loaded MSNPs. (b) Fluorescein (FITC) encapsulated MSNPs and (c) RhodamineB (RhoB) loaded MSNPs

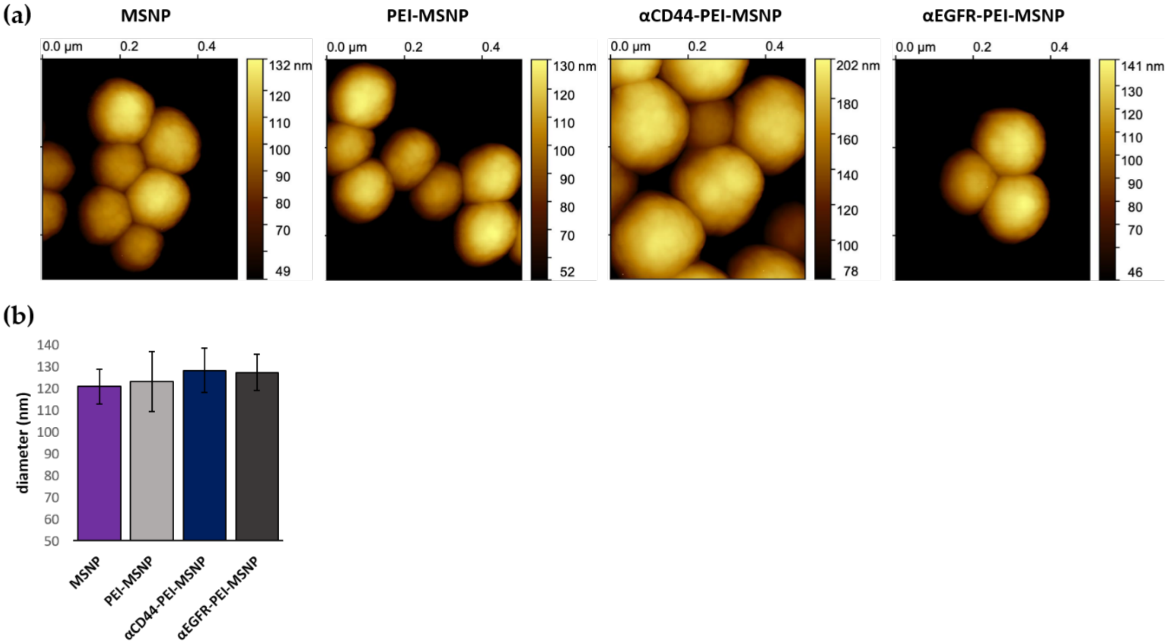


**Figure S2:** (a) AFM images of MSNPs, PEI-MSNP, $\alpha$CD44-PEI-MSNPs and $\alpha$CD44-PEI-MSNPs, (b) Graph of the average height measured with AFM (diameter). Error bars indicate ± SD.

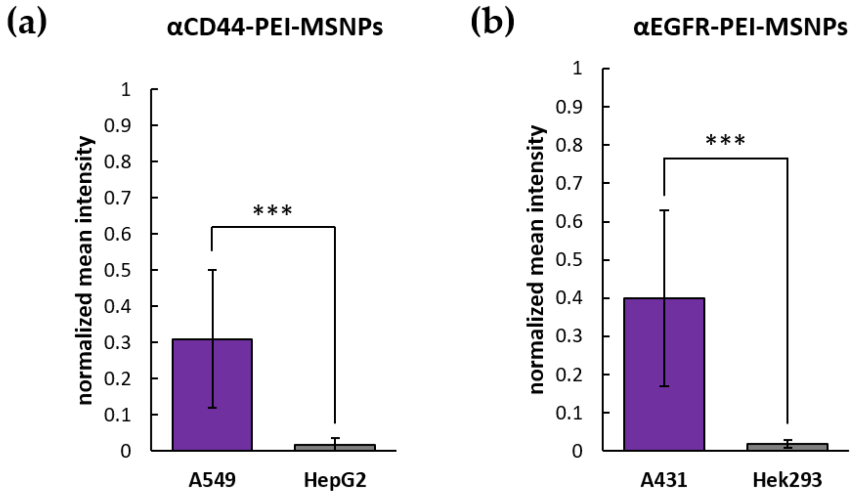


**Figure S3**: Quantitative analysis of the targeting efficiency of $\alpha$CD44-PEI-MSNPs and $\alpha$EGFR-PEI-MSNPs. (a) mean fluorescence intensity of $\alpha$CD44-PEI-MSNPs (encapsulated with RhoB) signal after 24 h incubation in A549 and HepG2 cells. (b) mean fluorescence intensity of $\alpha$EGFR-PEI-MSNPs (encapsulated with fluorescein) signal after 24 h incubation in A431 and Hek293 cells. Error bars indicate ± SD, with ***(p < 0,001).

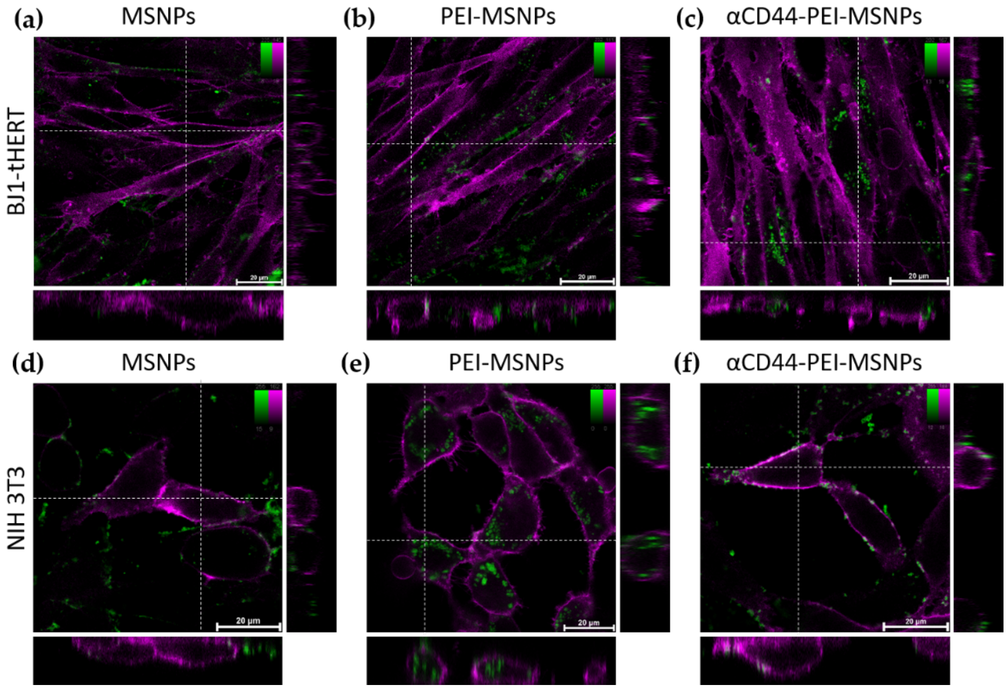


**Figure S4:** Confocal fluorescence microscopy images showing the influence of different MSNP coating in the uptake of nanoparticles in BJ1-tHERT and NIH 3T3 cells. Internalization of bare MSNP (a,d), PEI-coated MSNPs (PEI-MSNP, panels b,e) and CD44 functionalized MSNPs ($\alpha$CD44-PEI-MSNP, panels c,f) in BJ1-tHERT cells (a-c) and NIH 3T3 cells (d-f). Nanoparticles were loaded with RhoB (green) and the plasma membrane was stained with DiR (magenta). The central square represents a single xy plane, while the bottom and left panels are the xz and yz cross-sections, indicated by the dashed lines. Scale bar is 20 μM, color bars display the intensity values.

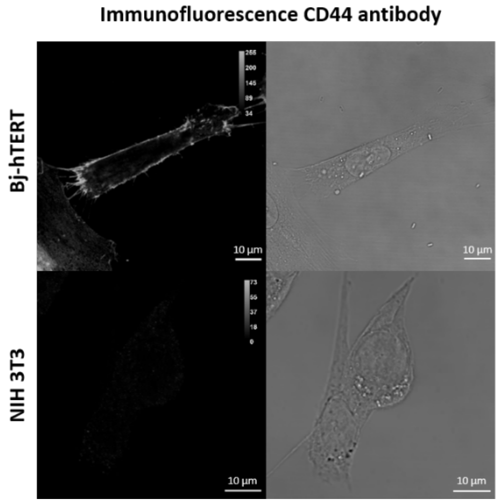


**Figure S5:** immunofluorescence (secondary goat-anti-rat IgG-AF488) staining showing the expression of absence of the CD44 receptor in Bj-hTERT and NIH 3T3, respectively. Scale bar is 10 µm.

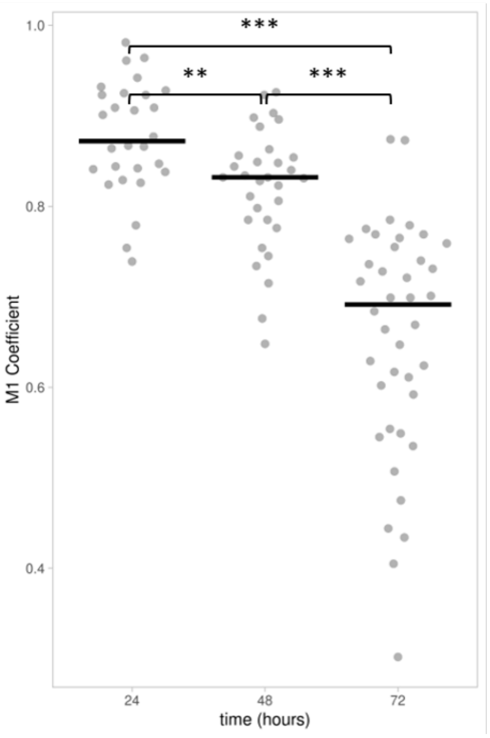


**Figure S6:** Manders' Coefficient (MC) representing the fraction of overlap between the $\alpha$CD44-PEI-MSNPs channel with the LysoTacker Deep Red channel. n values for 24, 48 and 72 h are 28, 31 and 40, respectively. ** ($p < 0{,}01$) and *** ($p < 0{,}001$)
